# Natural selection and recombination rate variation shape nucleotide polymorphism across the genomes of three related *Populus* species

**DOI:** 10.1101/026344

**Authors:** Jing Wang, Nathaniel R Street, Douglas G Scofield, Pär K Ingvarsson

**Affiliations:** Department of Ecology and Environmental Science, Umeå University, Umeå, SE 90187, Sweden; Umeå Plant Science Centre, Department of Plant Physiology, Umeå University, Umeå, SE 90187, Sweden; Department of Ecology and Genetics: Evolutionary Biology, Uppsala University, Uppsala, SE 75105, Sweden; Uppsala Multidisciplinary Center for Advanced Computational Science, Uppsala University, Uppsala, SE 75105, Sweden

**Keywords:** *Populus*, whole genome re-sequencing, nucleotide polymorphism, recombination, natural selection

## Abstract

A central aim of evolutionary genomics is to identify the relative roles that various evolutionary forces have played in generating and shaping genetic variation within and among species. Here we use whole-genome re-sequencing data to characterize and compare genome-wide patterns of nucleotide polymorphism, site frequency spectrum and population-scaled recombination rates in three species of *Populus*: *P. tremula, P. tremuloides* and *P. trichocarpa*. We find that *P. tremuloides* has the highest level of genome-wide variation, skewed allele frequencies and population-scaled recombination rates, whereas *P. trichocarpa* harbors the lowest. Our findings highlight multiple lines of evidence suggesting that natural selection, both due to purifying and positive selection, has widely shaped patterns of nucleotide polymorphism at linked neutral sites in all three species. Differences in effective population sizes and rates of recombination are largely explaining the disparate magnitudes and signatures of linked selection we observe among species. The present work provides the first phylogenetic comparative study at genome-wide scale in forest trees. This information will also improve our ability to understand how various evolutionary forces have interacted to influence genome evolution among related species.

## Introduction

A major goal in evolutionary genetics is to understand how genomic variation is established and maintained within and between species (Nordborg *et al*. 2005; Begun *et al*. 2007), and different evolutionary forces have substantial impacts in shaping genetic variation throughout the genome (Hellmann *et al*. 2005). Under the neutral theory, genetic variation is the manifestation of the balance between mutation and genetic drift (Kimura 1983). Demographic fluctuations, such as population expansion and/or bottlenecks, can cause patterns of genome-wide variation deviating from standard neutral model in various ways (Li and Durbin 2011). It is now clear, however, that natural selection, via positive selection favoring beneficial mutations (genetic hitchhiking) and/or purifying selection against deleterious mutations (background selection), plays an important role in moulding the landscape of nucleotide polymorphism in many species (Begun and Aquadro 1992; Begun *et al*. 2007; Cutter and Choi 2010; Mackay *et al*. 2012).

If natural selection is pervasive across the genome, patterns of genetic variation at linked neutral sites can be influenced by selection in a number of ways. First, positive correlations between levels of neutral polymorphism and recombination rates are expected since linked selection is expected to remove more neutral polymorphism in low-recombination regions compared to high-recombination regions and such a pattern is unlikely to be generated by demographic processes alone (Begun and Aquadro 1992; Kulathinal *et al*. 2008; McGaugh *et al*. 2012; Campos *et al*. 2014; Charlesworth and Campos 2014). Second, besides influencing the level of neutral variability, recombination rate can affect the efficacy of selection through the process known as Hill-Robertson interference (HBI) (Hill and Robertson 1966). If HRI is operating, genetic linkage effects in regions of low recombination will reduce the local effective population size (*N*_e_), and accordingly reduce the efficacy of selection (*N*_e_s), since the effects of selection are determined by the product of *N*_e_ and the selection coefficient on a mutation (s) (Kimura 1983). We would therefore expect both a reduced fixation of favorable mutations and an increased frequency of deleterious mutations in these regions (Hill and Robertson 1966; Haddrill *et al*. 2007; Campos *et al*. 2014). Third, signatures and magnitudes of linked selection are sensitive to the density of functional important sites (e.g. gene density) within specific genomic regions (Flowers *et al*. 2012). In accordance with the view that genes represent the most likely targets of natural selection, regions with a high density of genes are expected to have undergone stronger effects of linked selection and exhibit lower levels of neutral polymorphism (Nordborg *et al*. 2005; Flowers *et al*. 2012). Therefore, a positive or negative co-variation of recombination rate and gene density would act to either obscure or strengthen the signatures of linked selection across the genome (Cutter and Payseur 2003; Cutter and Choi 2010; Flowers *et al*. 2012). Lastly, a distinctive signature of recurrent selective sweeps is the local reduction of linked neutral polymorphism in regions experiencing frequent adaptive substitutions (Andolfatto 2007). A substantial number of adaptive substitutions are likely composed of amino acid substitutions and a negative correlation between neutral polymorphism and non-synonymous divergence can thus be particularly informative of the prevalence of selective sweeps (Macpherson *et al*. 2007). With the advance of next-generation sequencing technology, sufficient genome-wide data among multiple related species are becoming available (Luikart *et al*. 2003; Ellegren 2014). Phylogenetic comparative approaches will thus place us in a stronger position to understand how various evolutionary forces have interacted to shape the heterogeneous patterns of nucleotide polymorphism across the genome (Hufford *et al*. 2012; Cutter and Payseur 2013; Lawrie and Petrov 2014).

Thus far, genome-wide comparative studies have largely dealt with experimental model species, mammals, and cultivated plants of either agricultural or horticultural interest (Locke *et al*. 2011; Hufford *et al*. 2012; Liu *et al*. 2014). Forest trees, as a group, are characterized by extensive geographical distributions and are of high ecological and economic value (Neale and Kremer 2011). Most forest trees have largely persisted in an undomesticated state and, until quite recently, without anthropogenic influence (Neale and Kremer 2011). Accordingly, in contrast to crop and livestock lineages that have been through strong domestication bottlenecks, most extant populations of forest trees harbor a wealth of genetic variation and they are thus excellent model systems for dissecting the dominant evolutionary forces that sculpt patterns of variation throughout the genome (Gonzalez-Martinez *et al*. 2006; Neale and Kremer 2011). Among forest tree species, the genus *Populus* represents a particularly attractive choice because of its wide geographic distribution, important ecological role in a wide variety of habitats, multiple economic uses in wood and energy products, and relatively small genome size (Eckenwalder 1996; Jansson and Douglas 2007). Here, we studied three *Populus* species which differ in morphology, geographic distribution, population size and phylogenetic relationship (Figure S1) (Jansson *et al*. 2010; Wang *et al*. 2014). *P. tremula* and *P. tremuloides* (collectively ‘aspens’) have wide native ranges across Eurasia and North America respectively, are closely related, and belong to the same section of the genus (section *Populus*) (Jansson *et al*. 2010). In contrast, *P. trichocarpa* belongs to a different section of the genus (section *Tacamahaca*) that is reproductively isolated from members of the *Populus* section (Jansson *et al*. 2010). The distribution of *P. trichocarpa* is restricted to western regions of North America and its distribution range is considerably smaller than the two aspen species (Dickmann and Kuzovkina 2008). Importantly, *P. trichocarpa* also represents the first tree species to have its genome published (Tuskan *et al*. 2006) and the genome sequence and annotation have undergone continual improvement [http://phytozome.jgi.doe.gov]. This enables us to provide important context for our genome comparisons. The phylogenetic relationship of the three species ((P. *tremula-P. tremuloides*) *P. trichocarpa*) is well established by both chloroplast and nuclear DNA sequences (Hamzeh and Dayanandan 2004; Wang *et al*. 2014).

In this study, we used datasets generated by Next-Generation Sequencing (NGS) to characterize, compare and contrast genome-wide patterns of nucleotide diversity, site frequency spectrum, recombination rate, and to infer contextual patterns of selection throughout the genomes for all three species.

## Materials and Methods

### Samples and sequencing

Leaf samples were collected from 24 genotypes of *P. tremula* and 24 genotypes of *P. tremuloides* (Table S1). Genomic DNA was extracted from leaf samples, and paired-end sequencing libraries with insert sizes of 650bp were constructed for all genotypes. Whole-genome sequencing with a minimum expected depth of 20 × was performed on the Illumina HiSeq 2000 platform at the Science for Life Laboratory, Stockholm, Sweden and 2×100-bp paired-end reads were generated for all genotypes. Two samples of *P. tremuloides* failed to yield the expected coverage and were therefore removed from subsequent analyses. We obtained publicly available short read Illumina data of 24 *P. trichocarpa* individuals from NCBI SRA (Table S1). Individuals were selected to have a similar read depth as the samples of the two aspen species. The accession numbers of *P. trichocarpa* samples can be found in Evans *et al*. 2014. All analyses are thus based on data from 24 *P. tremula*, 22 *P. tremuloides* and 24 *P. trichocarpa* genotypes.

### Raw read filtering, read alignment and post-processing alignment

Prior to read alignment, we used Trimmomatic (Lohse *et al*. 2012) to remove adapter sequences from reads. Since the quality of reads always drops towards the end of reads, we used Trimmomatic to cut off bases from the start and/or end of reads when the quality values were smaller than 20. If the length of the processed reads was reduced to below 36 bases after trimming, reads were completely discarded. FastQC (http://www.bioinformatics.babraham.ac.uk/projects/fastqc/) was used to check and compare the per-base sequence quality between the raw sequence data and the filtered data. After quality control, all paired-end and orphaned single-end reads from each sample were mapped to the *P. trichocarpa* version 3 (v3.0) genome (Tuskan *et al*. 2006) using the BWA-MEM algorithm with default parameters in bwa-0.7.10 (Li 2013).

Several post-processing steps of alignments were performed to minimize the number of artifacts in downstream analysis: First, we performed indel realignment since mismatching bases were usually found in regions with insertions and deletions (indels) (Wang *et al*. 2015). The RealignerTargetCreator in GATK (The Genome Analysis Toolkit) (DePristo *et al*. 2011) was first used to find suspicious-looking intervals that were likely in need of realignment. Then, the IndelRealigner was used to run the realigner over those intervals. Second, as reads resulting from PCR duplicates can arise during the sequencing library preparation, we used the MarkDuplicates methods in the Picard package (http://picard.sourceforge.net) to remove those reads or read pairs having identical external coordinates and the same insert length. In such cases only the single read with the highest summed base qualities was kept for downstream analysis. Third, in order to exclude genotyping errors caused by paralogous or repetitive DNA sequences where reads were poorly mapped to the reference genome, or by other genome feature differences between *P. trichocarpa* and *P. tremula* or *P. tremuloides*, we removed sites with extremely low and extremely high read depths after investigating the empirical distribution of read coverage. We filtered out sites with a total coverage less than 100 × or greater than 1200 × across all samples per species. When reads were mapped to multiple locations in the genome, they were randomly assigned to one location with a mapping score of zero by BWA-MEM. In order to account for such misalignment effects, we removed those sites if there were more than 20 mapped reads with mapping score equaling to zero across all individuals in each species. Lastly, because the short read alignment is generally unreliable in highly repetitive genomic regions, we filtered out sites that overlapped with known repeat elements as identified by RepeatMasker (Tarailo-Graovac and Chen 2009). In the end, the subset of sites that passed all these filtering criteria in the three *Populus* species were used in downstream analyses.

### Single nucleotide polymorphism (SNP) and genotype calling

We implemented two complementary bioinformatics approaches: First, many studies have pointed out the bias inherent in population genetic estimates using genotype calling approach from NGS data (Nielsen *et al*. 2011; Nevado *et al*. 2014). Single- or multiple-sample genotype calling can result in a bias in the estimation of site frequency spectrum (SFS), as the former usually leads to overestimation of rare variants, whereas the latter often leads to the opposite (Nielsen *et al*. 2011). Therefore, in this study we employed a method, implemented in the software package - Analysis of Next-Generation Sequencing Data (ANGSD v0.602) (Korneliussen *et al*. 2014), to estimate the SFS and all population genetic statistics derived from the SFS without calling genotypes. Second, for those analyses that require accurate SNP and genotype calls, we performed SNP calling with HaplotypeCaller of the GATK v3.2.2 (DePristo *et al*. 2011), which called SNPs and indels simultaneously via local re-assembly of haplotypes for each individual and created single-sample gVCFs. GenotypeGVCFs in GATK was then used to merge multi-sample records together, correct genotype likelihoods, and re-genotype the newly merged record and perform re-annotation. Several filtering steps were then used to reduce the number of false positive SNPs and retain high-quality SNPs: (1) We removed all SNPs that overlapped with sites excluded by all previous filtering criteria. (2) We only retained bi-allelic SNPs with a distance of more than 5 bp away from any indels. (3) We treated genotypes with quality score (GQ) lower than 10 as missing and then removed those SNPs with genotype missing rate higher than 20%. (4) We removed SNPs that showed significant deviation from Hardy-Weinberg Equilibrium (P<0.001). After all filtering, 8,502,169 SNPs were detected among the three *Populus* species and were used in downstream analyses.

### Population structure

We used 4-fold synonymous SNPs with minor allele frequency >0.1 to perform population structure analyses with ADMIXTURE (Alexander *et al*. 2009). We ran ADMIXTURE on all the sampled individuals among species and on the samples within each species separately. The number of genetic clusters (*K*) was varied from 1 to 6. The most likely number of genetic cluster was selected by minimizing the cross-validation error in ADMIXTURE.

### Diversity and divergence - related summary statistics

For nucleotide diversity and divergence estimates, only the reads with mapping quality above 30 and the bases with quality score higher than 20 were used in all downstream analyses with ANGSD (Korneliussen *et al*. 2014). First, we used the -doSaf implementation in ANGSD to calculate the site allele frequency likelihood based on the SAMTools genotype likelihood model (Li *et al*. 2009). Then, we used the -realSFS implementation in ANGSD to obtain an optimized folded global SFS using Expectation Maximization (EM) algorithm for each species. Based on the global SFS, we used the -doThetas function in ANGSD to estimate the per-site nucleotide diversity from posterior probability of allele frequency based on a maximum likelihood approach (Kim *et al*. 2011). Two standard estimates of nucleotide diversity, the average pairwise nucleotide diversity (Θ_π_) (Tajima 1989) and the proportion of segregating sites (Θ_W_) (Watterson 1975), and one neutrality statistic test Tajima’s D (Tajima 1989) were summarized along all 19 chromosomes using non-overlapping sliding windows of 100 kilobases (Kbp) and 1 megabases (Mbp). Windows with less than 10% of covered sites left from previous quality filtering steps were excluded. In the end, 3340 100-Kbp and 343 1-Mbp windows, with an average of 50,538 and 455,910 covered bases per window, were respectively included.

All these statistics were also calculated for each type of functional element (0fold non-synonymous, 4-fold synonymous, intron, 3’ UTR, 5’ UTR, and intergenic sites) over non-overlapping 100-Kbp and 1-Mbp windows in all three *Populus* species. The category of gene models followed the gene annotation of *P. trichocarpa* version 3.0 (Tuskan *et al*. 2006). For protein-coding genes, we only included genes with at least 90% of covered sites left from previous filtering steps to ensure that the three species have the same gene structures. For regions overlapped by different transcripts in each gene, we classified each site according to the following hierarchy (from highest to lowest): Coding regions (CDS), 3’UTR, 5’UTR, Intron. Thus, if a site resides in a 3’UTR in one transcript and CDS for another, the site was classified as CDS. We used the transcript with the highest content of protein-coding sites to categorize synonymous and non-synonymous sites within each gene. A respective of 16.52 Mbp, 3.4 Mbp, 7.19 Mbp, 4.02 Mbp, 31.89 Mbp and 73.46 Mbp were partitioned into 0-fold non-synonymous (where all DNA sequence changes lead to protein sequence changes), 4-fold synonymous (where all DNA sequence changes lead to the same protein sequences), 3’UTR, 5’UTR, intron, and intergenic categories.

### Linkage disequilibrium (LD) and population-scaled recombination rate (*ρ*)

A total of 1,409,377 SNPs, 1,263,661 SNPs and 710,332 SNPs with minor allele frequency higher than 10% were used for the analysis of LD and *ρ* in *P. tremula, P. tremuloides* and *P. trichocarpa*, respectively. To estimate and compare the rate of LD decay among the three *Populus* species, we firstly used PLINK 1.9 (Purcell *et al*. 2007) to randomly thin the number of SNPs to 100,000 in each species. We then calculated the squared correlation coefficients (*r*^2^) between all pairs of SNPs that were within a distance of 50 Kbp using PLINK 1.9. The decay of LD against physical distance was estimated using nonlinear regression of pairwise *r*^2^ vs. the physical distance between sites in base pairs (Remington *et al*. 2001).

We estimated the population-scaled recombination rate *ρ* using the Interval program of LDhat 2.2 (McVean *et al*. 2004) with 1,000,000 MCMC iterations sampling every 2,000 iterations and a block penalty parameter of five. The first 100,000 iterations of the MCMC iterations were discarded as a burn-in. We then calculated the scaled value of *ρ* in each 100-Kbp and 1-Mbp window by averaging over all SNPs in that window. Only windows with more than 10,000 (in 100 Kbp windows) and 100,000 sites (in 1 Mbp windows) and 100 SNPs left from previous filtering steps were used for the estimation of *ρ*.

### Estimating the distribution of fitness effects of new amino acid mutations (DFE) and the proportion of adaptive amino acid substitutions (α)

We generated the folded SFS in each species for a class of selected sites (0-fold non-synonymous sites) and a class of putatively neutral reference sites (4-fold synonymous sites) from SNPs data using a custom Perl script. We employed a maximum likelihood (ML) approach as implemented in the program DFE-alpha (Keightley and Eyre-Walker 2007; Eyre-Walker and Keightley 2009) to fit a demographic model with a step of population size change to the neutral SFS. Fitness effects of new deleterious mutations at the selected site class were sampled from a gamma distribution after incorporating the estimated parameters for the demographic model. This method assumes that fitness effects of new mutations at neutral sites are zero and unconditionally deleterious at selected sites since it assumes that advantageous mutations are too rare to contribute to polymorphism (Keightley and Eyre-Walker 2007). We report the proportion of amino acid mutations falling into different effective strengths of selection (*N*_e_s) range: 0-1, 1-10, >10, respectively.

From the estimated DFE, the proportion (α) and the relative rate (ω) of adaptive substitution at 0-fold non-synonymous sites were estimated using the method of Eyre-Walker and Keightley (2009). This method explicitly account for past changes in population size and the presence of slightly deleterious mutations. Among the total of 8,502,169 SNPs detected by GATK, on average less than 1% were shared between either of the two aspen species and *P. trichocarpa* (Figure S2). We therefore used the aspen species and *P. trichocarpa* as each other’s outgroup species to calculate between-species nucleotide divergence at 4-fold synonymous and 0-fold non-synonymous sites since it is unlikely to be influenced by shared ancestral polymorphisms. Jukes-Cantor multiple hits correction was applied to the divergence estimates (Jukes and Cantor 1969). For the parameters of *N*_e_s, α and ω, we generated 200 bootstrap replicates by resampling randomly across all SNPs in each site class using R (R Develpment Core Team 2014). We excluded the top and bottom 2.5% of bootstrap replicates and used the remainder to represent the 95% confidence intervals for each parameter.

### Genomic correlates of diversity

In order to examine the factors influencing levels of neutral polymorphism in all three *Populus* species, we firstly assume that the 4-fold synonymous sites in genic regions were selectively neutral as every possible mutation at 4-fold degenerate sites is synonymous. In the following we refer to the pairwise nucleotide diversity at 4-fold synonymous sites (θ_4-fold_) as “neutral polymorphism”. As a comparison to genic region, we also estimated levels of nucleotide diversity at intergenic sites (θ_Intergenic_). Then, we tabulated several other genomic features within each 100 Kbp and 1 Mbp window that may correlate with patterns of polymorphism. First, we summarized population-scaled recombination rate (ρ) as described above for each species. Second, we tabulated GC content as the fraction of bases where the reference sequence *(P. trichocarpa* v3.0) was a G or a C. Third, we measured the gene density as the number of functional genes within each window according to the gene annotation of *P. trichocarpa* version 3.0. Any portion of a gene that fell within a window was counted as a full gene. Fourth, we accounted for the variation of mutation rate by calculating the number of fixed differences per neutral site (either 4-fold synonymous sites or intergenic sites) between aspen and *P. trichocarpa* within each window, which was performed in the ngsTools (Fumagalli *et al*. 2014). The reason why we used divergence between aspen and *P. trichocarpa* to measure mutation rate is because they are distantly related (Wang *et al*. 20141), and the estimate of divergence are unlikely to be influenced by shared ancestral polymorphisms between species as shown above. Fifth, we tabulated the number of covered bases in each window as those left from all previous filtering criteria.

We used Spearman’s rank-order correlation test to examine pairwise correlations between the variables of interest. In order to account for the autocorrelation between variables, we further calculated partial correlations between the variables of interest by removing the confounding effects of other variables (Kim and Soojin 2007). All statistical tests were performed using R version 3.2.0 unless stated otherwise.

### Data Availability

All newly generated Illumina reads of 24 *P. tremula* and 22 *P. tremuloides* from this study have been submitted to the Short Read Archive (SRA) at NCBI. All accession IDs can be found in Table S1.

## Results

We generated whole-genome sequencing data for 24 *P. tremula* and 22 *P. tremuloides* (Table S1) with all samples sequenced to relatively high depth (24.2×-69.2×; Table S2). We also downloaded whole-genome re-sequencing data for 24 samples of *P. trichocarpa* from the NCBI Short Read Archive (Evans et al. 2014). After adapter removal and quality trimming, 949.2 Gb of high quality sequence data remained (Figure S3; Table S2). The mean mapping rate of reads to the *P. trichocarpa* reference genome were 89.8% for *P. tremula*, 91.1% for *P. tremuloides*, and 95.2% for *P. trichocarpa* (Table S2). On average, the genome-wide coverage of uniquely mapped reads was more than 20 × for each species (Table S2). After excluding sites with extreme coverage, low mapping quality, or those overlapping with annotated repetitive elements (see Materials and Methods), 42.8% of collinear genomic sequences remained for downstream analyses. 54.9% of these sites were found within gene boundaries, covering 70.1% of all genic regions predicted from the *P. trichocarpa* assembly. The remainder sites (45.1%) were located in intergenic regions.

### Genome-wide patterns of polymorphism, site frequency spectrum and recombination among the three Populus species

When analyzing population structure between species, we found the model exhibited the lowest cross-validation error when the number of ancestral populations (*K*) = 3 (Figure S4b), which clearly subdivided the three species into three distinct clusters (Figure S4a). When we analyzed population structure within each species separately, we found that the cross-validation error increased linearly with increasing *K*, with *K* = 1 minimizing the cross-validation error for all three species (Figure S4c-e). Our results are slightly different from several earlier studies that have documented population subdivision in these species (De Carvalho *et al*. 2010; Evans *et al*. 2014) but it is likely due to the small sample sizes used in this study (22-24 individuals). In addition, it could also be caused by the low power of model-based approaches in detecting population structure when it is very weak (Alexander *et al*. 2009). More generally, population structure is expected to be weak in *Populus* given the great dispersal capabilities of both pollen and seeds (Eckenwalder 1996; Jansson and Douglas 2007).

The two aspen species harbor substantial levels of nucleotide diversity across the genome (Θп=0.0133 in *P. tremula*; Θп=0.0144 in *P. tremuloides*), approximately two to three-fold higher than diversity in *P. trichocarpa* (Θп=0.0059) (Figure 1; Table S3). Among various genomic contexts, we found the levels of nucleotide diversity were highest at intergenic sites, followed by 4-fold synonymous sites, 3’UTRs, 5’UTRs, introns and were lowest at 0-fold non-synonymous sites (Figure S5; Table S3). In accordance with the view that the large majority of amino acid mutations are selected against (Larracuente *et al*. 2008), we found significantly lower Tajima’s D at 0-fold non-synonymous sites compared to 4-fold synonymous sites (P<0.001, Mann-Whitney U test) (Figure S6; Table S3). In addition, we observed significantly positive correlations of Θп between each pair of the three species across the whole genome (Figure 2a). The overall nucleotide diversity estimated in *P. trichocarpa* was slightly higher than the value reported in Evans *et al*. 2014 (Θп=0.0041), but this likely only reflects differences between the methods used in the two studies. In this study, we utilized the full information of the filtered data and estimated the population genetic statistics directly from genotype likelihoods, which take statistical uncertainty of SNP and genotype calling into account and should give more accurate estimates (Kim *et al*. 2011; Nielsen *et al*. 2011).

**Figure 1.**
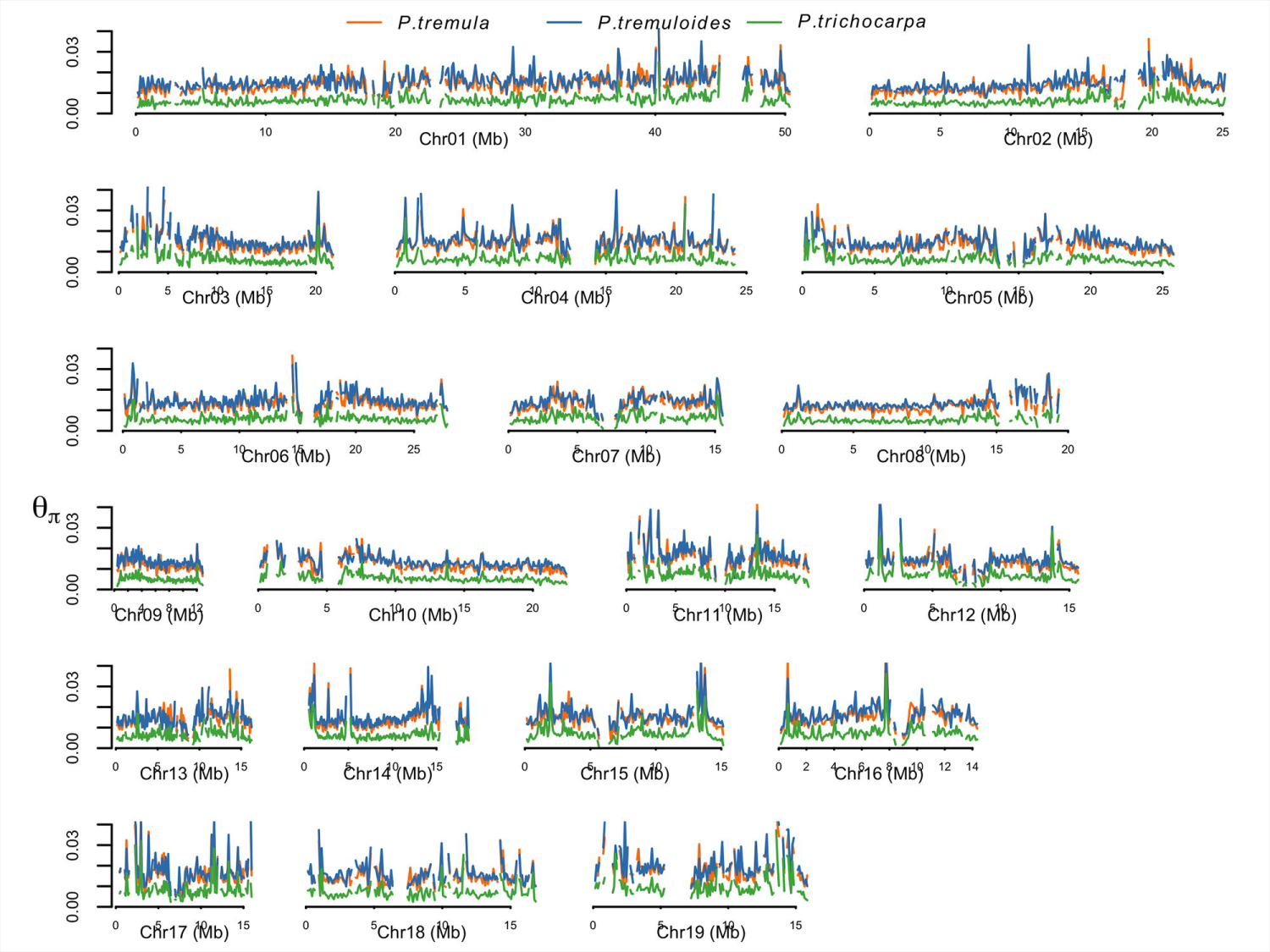
Genome-wide patterns of polymorphism among three *Populus* species. Nucleotide diversity (Θ_π_) was calculated over 100 Kbp non-overlapping windows in *P. tremula* (orange line), *P. tremuloides* (blue line) and *P. trichocarpa* (green line) along the 19 chromosomes.

**Figure 2.**
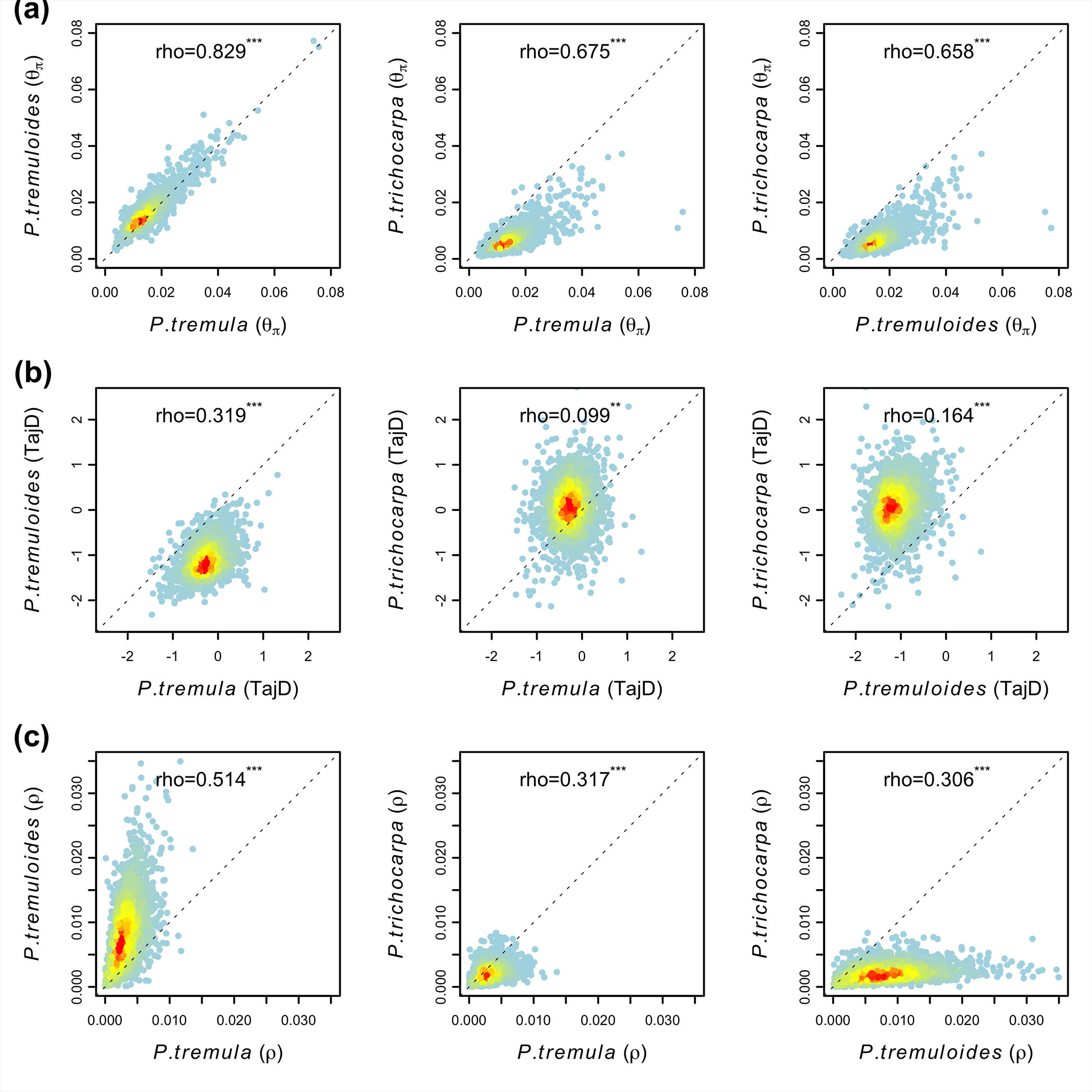
Distribution and correlations of (a) polymorphism (Θ_π_), (b) Tajima’s D and (c) population-scaled recombination rate (ρ) between pairwise comparisons of *P. tremula, P. tremuloides* and *P. trichocarpa* over 100 Kbp non-overlapping windows. The red to yellow to blue gradient indicates decreased density of observed events at a given location in the graph. Spearman’s rank correlation coefficient (rho) and the *P*-value are shown in each subplot. (*** *P*<2.2×10^-16^, \*\**P*<0.001). The dotted grey line in each subplot indicates simple linear regression line with intercept being zero and slope being one.

Compared to patterns of polymorphism, we observed much weaker correlations of the site frequency spectrum, summarized using the Tajima’s D statistic (Tajima 1989), between species (Figure 2b). *P. tremuloides* (average Tajima’s D=-1.169) showed substantially greater negative values of Tajima’s D along all chromosomes compared to both *P. trichocarpa* (average Tajima’s D=0.064) and *P. tremula* (average Tajima’s D=-0.272) (Figure S7; Table S3), reflecting a large excess of low-frequency polymorphisms segregating in this species. Furthermore, the three *Populus* species showed different extents of genome-wide LD decay (Figure S8), with LD decaying fastest in *P. tremuloides* and slowest in *P. trichocarpa* (Figure S8). This reflects the rank order of their population-scaled recombination rates (*ρ*=4*N*_e_c) (Figure S9), for which the mean *ρ* over 100 Kbp non-overlapping windows was highest in *P. tremuloides* (8.42 Kbp^-1^), followed by *P. tremula* (3.23 Kbp^-1^), and lowest in *P. trichocarpa* (2.19 Kbp^-1^). Intermediate correlations of recombination rates were observed between species (Figure 2c). In addition, concordant values of Θ_п_, Tajima’s D and *ρ* for all three species were also observed in 1 Mbp windows (Figure S10). For populations under drift-mutation-recombination equilibrium, *ρ* = 4*N*_e_c (where *N*_e_ is the effective population size and *c* is the recombination rate) and Θ_W_ = 4*N*_e_*μ* (where *N*_e_ is the effective population size and *μ* is the mutation rate). In order to compare the relative contribution of recombination (*c*) and mutation (*μ*) in shaping genomic variation, we measured the ratio of population recombination rate to the nucleotide diversity (*ρ*/θ_W_) across the genome (Figure S11). The mean *c*/*μ* in *P. tremula, P. tremuloides* and *P. trichocarpa* was 0.22, 0.39 and 0.38 respectively.

### Distribution of fitness effects and proportion of adaptive amino acid substitutions

We quantified the efficacy of both purifying and positive selection using the information of polymorphism and divergence among the three species. The estimated distribution of fitness effects of new 0-fold non-synonymous mutations indicate that the majority of new amino acid mutations were strongly deleterious (*N*_e_s>10) and likely to be under strong levels of purifying selection in all three species (Figure 3; Table S4). There was a greater proportion of amino acid mutations under moderate levels of purifying selection (1<*N*_e_s<10) in *P. tremuloides* (∼31%), compared to *P. tremula* (∼16%) and *P. trichocarpa* (∼10%). In comparison, we found a higher proportion of weakly deleterious mutations that behave as effectively neutral (*N*_e_s<1) in *P. trichocarpa* (∼31%) relative to *P. tremula* (∼23%) and *P. tremuloides* (∼16%) (Figure 3; Table S4).

**Figure 3.**
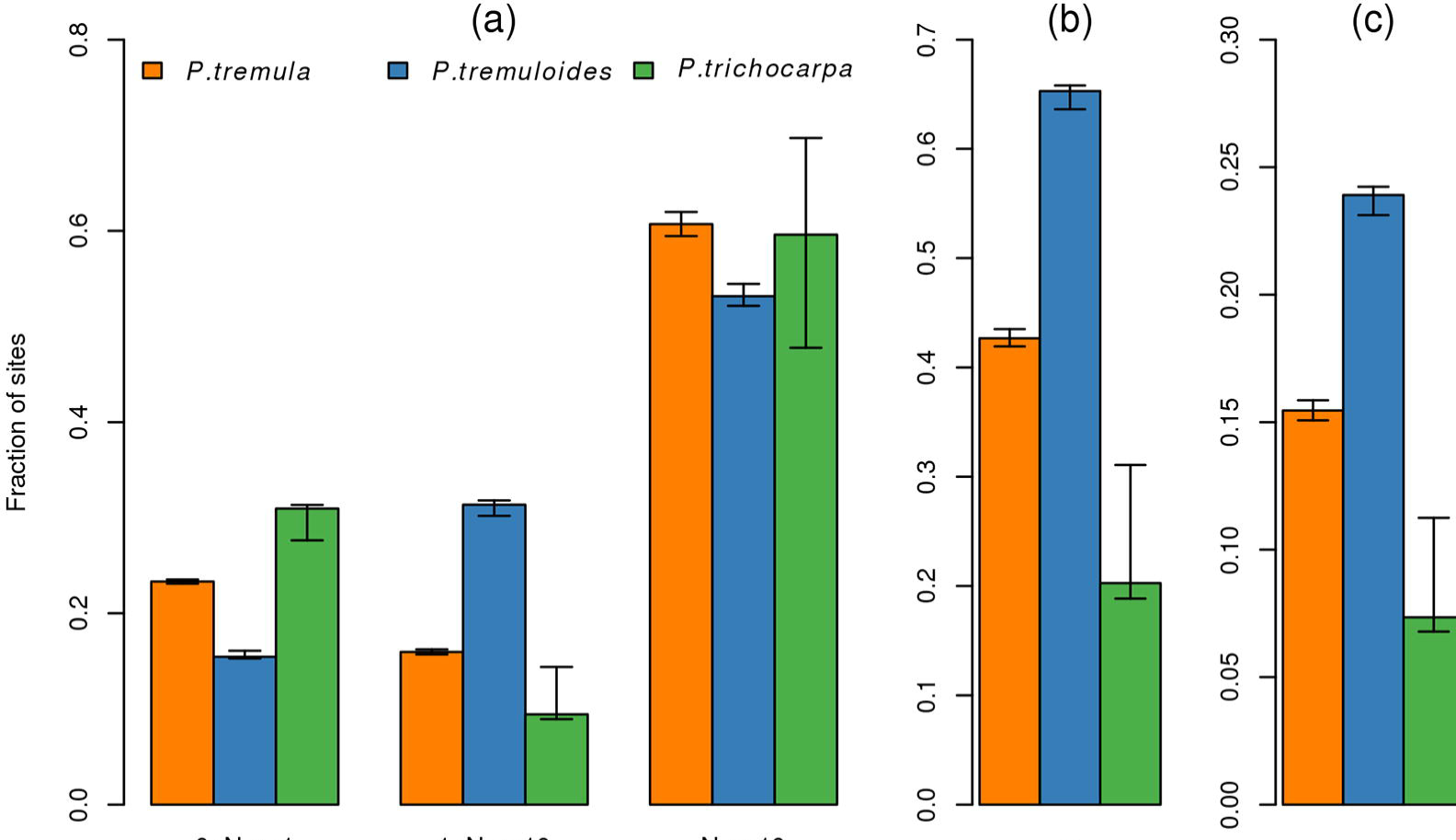
Estimates of purifying and positive selection at 0-fold non-synonymous sites in three *Populus* species. (a) The distribution of fitness effects of new amino acid mutations (DFE), (b) the proportion of adaptive substitution (α), and (c) the rate of adaptive non-synonymous to synonymous substitutions (ω) in *P. tremula* (orange bar), *P. tremuloides* (blue bar) and *P. trichocarpa* (green bar). Error bars represent 95% bootstrap confidence intervals.

Using 4-fold synonymous sites as a neutral reference, we employed an extension of the McDonald-Kreitman test (Eyre-Walker and Keightley 2009) to estimate the fraction of adaptive amino acid substitutions (α) and the rate of adaptive substitution relative to the rate of neutral substitution (ω) in all three species. Both α and ω were highest in *P. tremuloides* (α: ∼65% [95% CI: 63.6%-65.8%]; ω: ∼0.24 [95% CI: 0.231-0.242]), intermediate in *P. tremula* (α: ∼43% [95% CI: 41.9%-43.5%]; ω: ∼0.16 [95% CI: 0.151-0.159]) and lowest in *P. trichocarpa* (α: ∼20%[95% CI: 18.8%-31.1%]; ω: ∼0.07 [95% CI: 0.068-0.112]) (Figure 3; Table S4).

### Neutral polymorphism, but not divergence, is positively correlated with recombination rate

If natural selection (either purifying or positive selection) occurs throughout the genome at similar rates, they should leave a stronger imprint on patterns of neutral polymorphism in regions experiencing low recombination (Begun and Aquadro 1992). In accordance with this expectation, we found significantly positive correlations between levels of neutral polymorphism (θ_4-fold_) and population recombination rates in both aspen species (Table 1), with correlations being stronger in *P. tremula* than in *P. tremuloides*. In *P. trichocarpa*, however, we found either no or weak correlation between diversity and recombination rate (Table 1). Compared to 100 Kbp windows, correlations were stronger for 1 Mbp windows in all species, which most likely results from the higher signal-to-noise ratio provided by larger genomic windows (Table 1). In the remainder of this paper we thus focus our analyses primarily on data generated with a 1 Mbp window size. When performing simple linear regression analysis between diversity and recombination rate over 1 Mbp windows, the recombination rate explained 45.8%, 21.3%, and 3.9% of the amount of neutral genetic variation in *P. tremula, P. tremuloides* and *P. trichocarpa*, respectively (Figure 4). If the positive relationship between diversity and recombination rate was merely caused by the mutagenic effect of recombination, similar patterns should also be observed between divergence and recombination rate (Kulathinal *et al*. 2008). However, no such correlations were observed in any of the three species (Table 1; Figure 4). The correlations between neutral diversity and recombination rate were slightly lower, but still significant, after using partial correlations to control for possible confounding factors such as GC content, gene density, divergence at neutral sites, and the number of neutral bases covered by sequencing data (Table 1).

**Table 1.** Summary of the correlation coefficients (Spearman’s rank correlation coefficient) between levels of neutral polymorphism (Θ), divergence (d) and recombination rate (*ρ*) in genic and intergenic regions among all three *Populus* species.

| Dataset | Species | $\rho$ vs. $\Theta_{4\text{-fold}}$ | | $\rho$ vs. $d_{4\text{-fold}}$ | $\rho$ vs. $\Theta_{\text{Intergenic}}$ | | $\rho$ vs. $d_{\text{Intergenic}}$ |
| --- | --- | --- | --- | --- | --- | --- | --- |
|  |  | Pairwise | Partial<sup>a</sup> |  | Pairwise | Partial<sup>b</sup> |  |
| 100Kbp | *P. tremula* | 0.339<sup>***</sup> | 0.309<sup>***</sup> | 0.043 | 0.062<sup>**</sup> | 0.142<sup>***</sup> | -0.077<sup>**</sup> |
|  | *P. tremuloides* | 0.310<sup>***</sup> | 0.284<sup>***</sup> | 0.061<sup>**</sup> | -0.037 | 0.100<sup>**</sup> | -0.029 |
|  | *P. trichocarpa* | 0.011 | -0.024 | 0.053<sup>*</sup> | -0.080<sup>**</sup> | -0.002 | -0.015 |
| 1Mbp | *P. tremula* | 0.647<sup>***</sup> | 0.573<sup>***</sup> | -0.070 | 0.201<sup>**</sup> | 0.348<sup>**</sup> | -0.209<sup>**</sup> |
|  | *P. tremuloides* | 0.400<sup>**</sup> | 0.363<sup>**</sup> | -0.033 | 0.032 | 0.320<sup>**</sup> | -0.127<sup>*</sup> |
|  | *P. trichocarpa* | 0.227<sup>**</sup> | 0.151<sup>*</sup> | -0.027 | -0.072 | 0.165<sup>*</sup> | -0.120<sup>*</sup> |

<sup>a</sup>Partial correlation controls for GC content, gene density, divergence of 4-fold synonymous sites between aspen and *P. trichocarpa*, and coverage (the number of 4-fold synonymous bases covered by sequencing data).
<sup>b</sup>Partial correlation controls for GC content, gene density, divergence of intergenic sites between aspen and *P. trichocarpa*, and coverage (the number of intergenic bases covered by sequencing data).
\* $P < 0.05$
\*\* $P < 0.001$
\*\*\* $P < 2.2 \times 10^{-16}$

**Figure 4.**
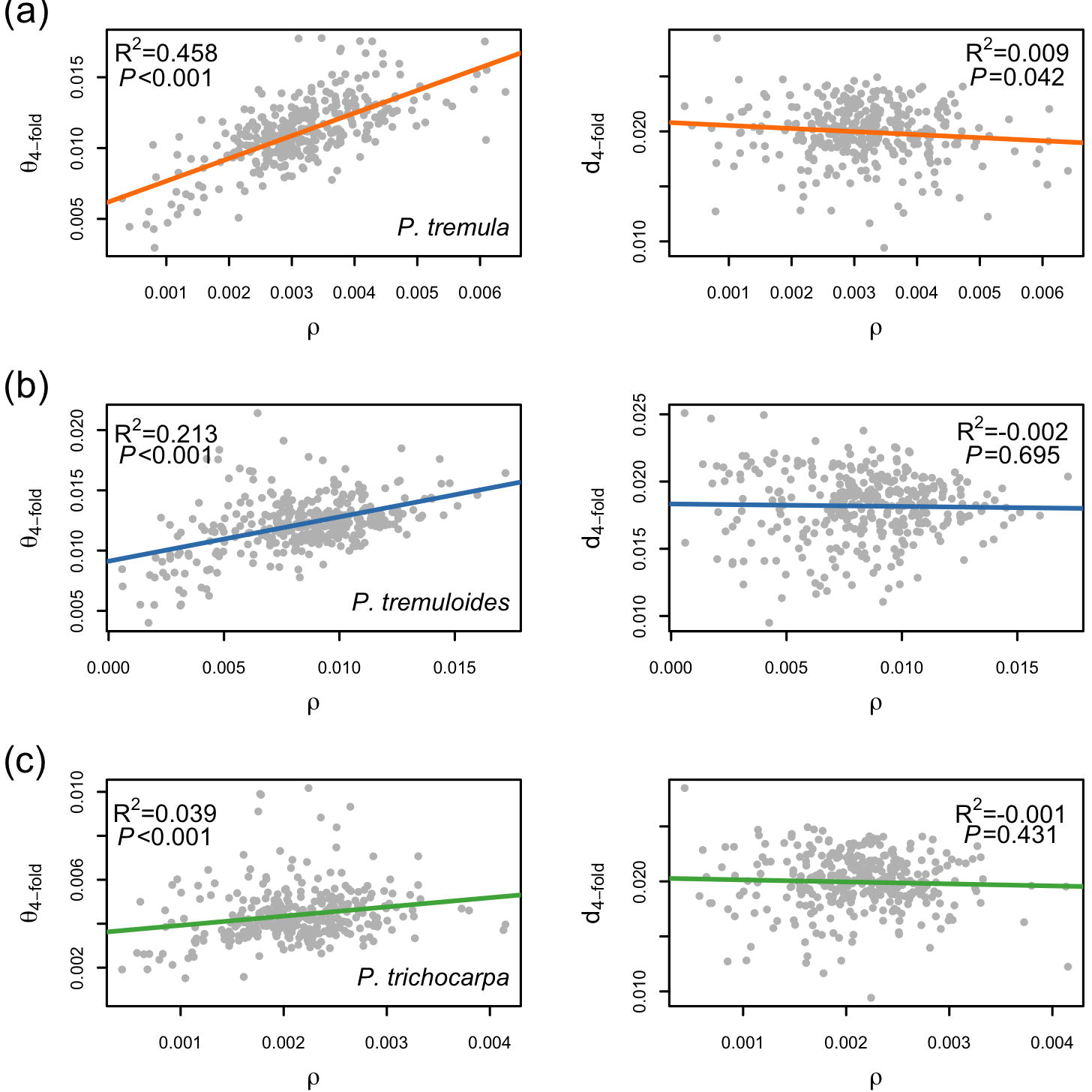
Correlations of estimates between neutral genetic diversity (Θ_4-fold_) (left panel), neutral genetic divergence (d_4-fold_) (right panel) and population-scaled recombination rates (*ρ*) over 1Mbp non-overlapping windows. Linear regression lines are colored according to species: (a) *P. tremula* (orange line), (b) *P. tremuloides* (blue line) and (c) *P. trichocarpa* (green line).

In accordance with the view that genes represent the most likely targets of natural selection (Lohmueller *et al*. 2011), the correlations between intergenic diversity and recombination rate were substantially weaker than those correlations in genic regions (Table 1). Only 7.3% of the variation in intergenic nucleotide diversity in *P. tremula* could be explained by variation in the recombination rate, whereas the impact of recombination rate variation on intergenic diversity in *P. tremuloides* and *P. trichocarpa* was negligible (<1% Figure S12; Table 1). However, after using partial correlation analyses to control for possible confounding factors, the correlations between intergenic diversity and recombination rate became significant in all species. Compared to genic regions these correlations were slightly higher in *P. trichocarpa*, of similar magnitude in *P. tremuloides* and weak in *P. tremula* (Table 1).

### The effect of recombination on the efficacy of natural selection

We characterized the ratio of non-synonymous to synonymous polymorphism (θ_0-fold_/θ_4-fold_) and divergence (d_0-fold_/d_4-fold_) to assess whether there was a relationship between the efficacy of natural selection and the rate of recombination (Table 2). Once GC content, gene density and the number of 4-fold synonymous and 0-fold non-synonymous sites were taken into account, we found no correlation between recombination rate and d_0-fold_/d_4-fold_ in any of the three species (Table 2). We also did not observe any significant correlations between recombination rate and θ_0-fold_/θ_4-fold_ over 1 Mbp windows after controlling for confounding factors (Table 2). However, when using 100 Kbp windows, we found significantly negative correlations between recombination rate and θ_0-fold_/θ_4-fold_ in *P. tremula* and *P. tremuloides*, but not in *P. trichocarpa* (Table 2).

**Table 2.**
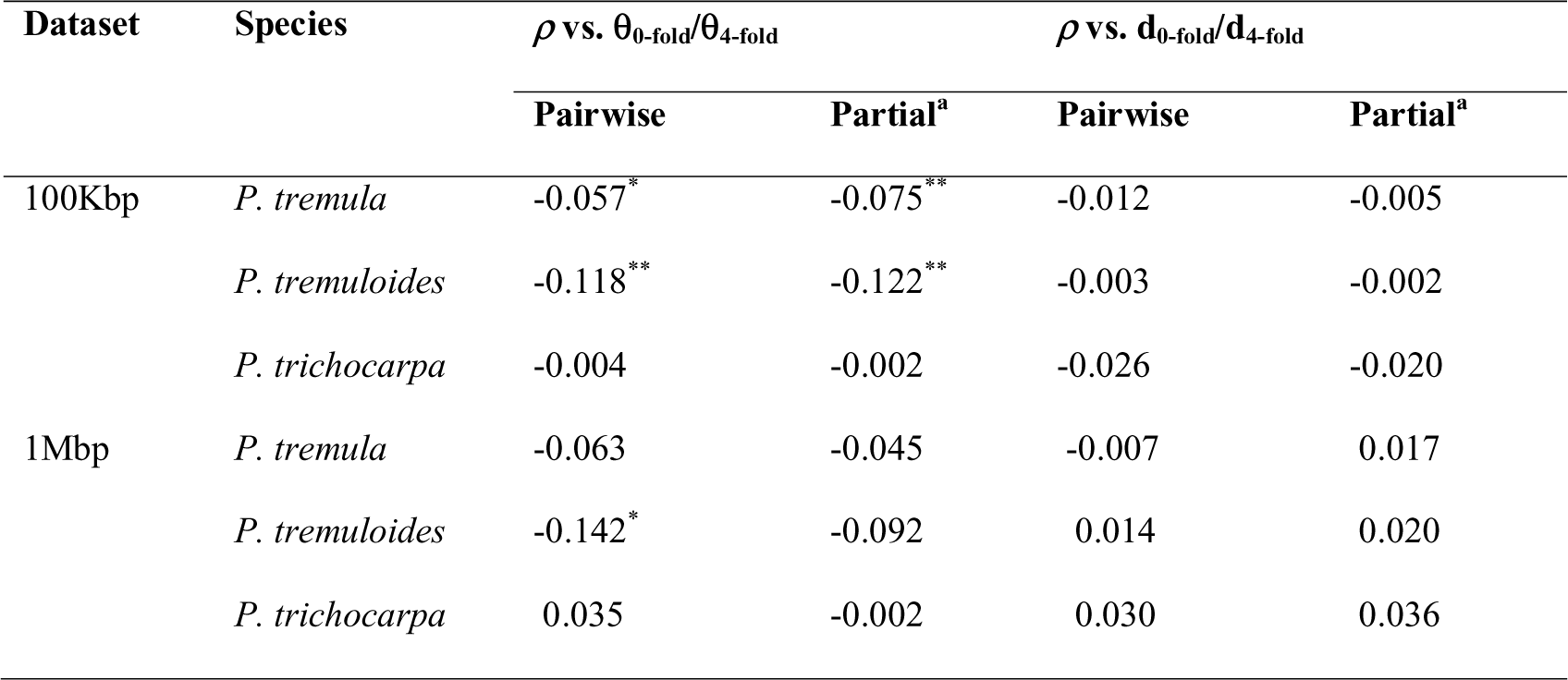
Summary of the correlation coefficients (Spearman’s rank correlation coefficient) between recombination rate (*ρ*) and the ratio of non-synonymous to synonymous polymorphism (θ_0-fold_/θ_4-fold_) and divergence (d_0-fold_/d_4-fold_).

### Inconsistent effect of gene density on patterns of polymorphism in genic vs. intergenic regions

We measured gene density as the number of protein-coding genes in each 1 Mbp window, which in turn was highly correlated with the proportion of coding bases in each window (Figure S13). For all three species, we found significantly positive correlations between population recombination rate and gene density (Figure 5a; Table 3). However, rather than being linear, the relationships between recombination rate and gene density was curvilinear with a significant positive correlation observed only in regions of low gene density (gene number smaller than ∼85 within each 1Mbp window) (Table 3). For regions of high gene density (gene number greater than ∼85 within each 1Mbp window) we found no correlations between recombination rate and gene density in both aspen species, and only a weak, positive correlation in *P. trichocarpa* (Figure 5a; Table 3). After controlling for GC content and the number of bases covered by sequencing data, the correlation became significant in regions of high gene density for *P. tremula*, but remained non-significant for *P. tremuloides* (Table 3).

**Figure 5.**
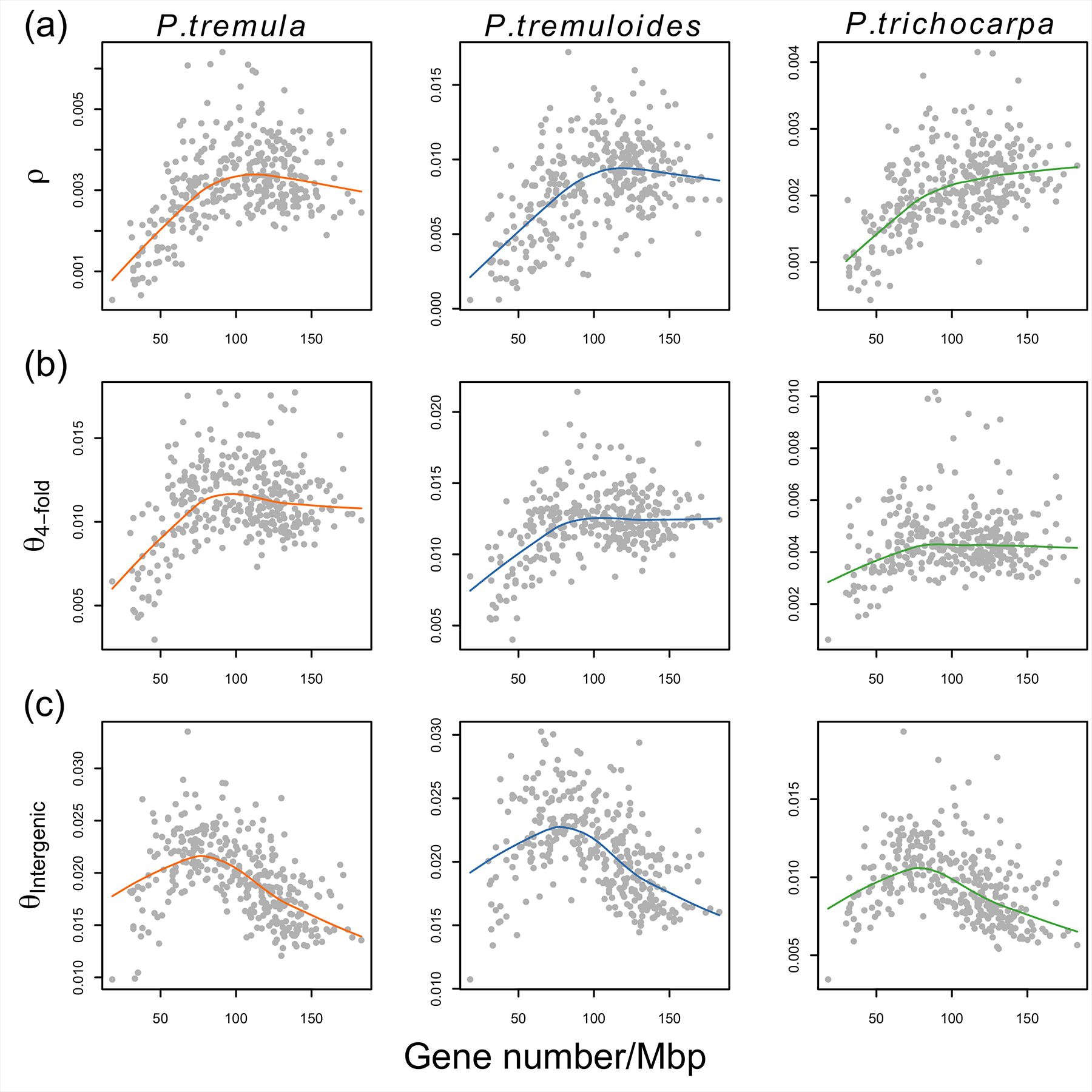
Correlations of estimates between (a) population-scaled recombination rates (*ρ*), (b) genic genetic diversity (Θ_4-fold_), (c) intergenic genetic diversity (Θ_Intergenic_) and gene density over 1 Mbp non-overlapping windows in *P. tremula* (left panel), *P. tremuloides* (middle panel) and *P. trichocarpa* (right panel). Grey points represent the statistics computed over 1Mbp non-overlapping windows. Colored lines denote the lowess curves fit to the analyzed two variables in each species.

**Table 3.** Summary of the correlation coefficients (Spearman’s rank correlation coefficient) between gene density and population recombination rate (*ρ*), neutral polymorphism in genic (Θ_4-fold_) and intergenic regions (Θ_intergenic_) over 1 Mbp non-overlapping windows in three *Populus* species.

| Species | Correlation type | Gene density vs. $\rho^a$ | | Gene density vs. $\Theta_{4\text{-fold}}^b$ | | Gene density vs. $\Theta_{\text{intergenic}}^c$ | |
| --- | --- | --- | --- | --- | --- | --- | --- |
|  |  | low | high | low | high | low | high |
| *P. tremula* | Pairwise | 0.674** | -0.112 | 0.601** | -0.180* | 0.431** | -0.605*** |
|  | Partial | 0.516** | 0.263* | 0.191* | 0.110 | 0.263* | -0.438** |
| *P.tremuloides* | Pairwise | 0.527** | 0.006 | 0.576** | -0.077 | 0.419** | -0.600*** |
|  | Partial | 0.315** | 0.048 | 0.407** | 0.280** | 0.363** | -0.444** |
| *P.trichocarpa* | Pairwise | 0.609** | 0.168* | 0.417** | -0.033 | 0.529** | -0.513*** |
|  | Partial | 0.477** | 0.193* | 0.242* | 0.263** | 0.432** | -0.273** |

<sup>a</sup>Partial correlation controls for GC content and the number of bases covered by the data
<sup>b</sup>Partial correlation controls for GC content, population recombination rate, divergence of 4-fold synonymous sites between aspen and *P. trichocarpa*, and coverage (the number of 4-fold synonymous bases covered by sequencing data).
<sup>c</sup>Partial correlation controls for GC content, population recombination rate, divergence of intergenic sites between aspen and *P. trichocarpa*, and coverage (the number of intergenic bases covered by sequencing data).
\* $P < 0.05$
\*\* $P < 0.001$

We then examined the relationship between neutral polymorphism and gene density. Compared to the prediction of lower diversity in regions with higher functional density (Payseur and Nachman 2002), we found that the correlation pattern between gene density and levels of neutral polymorphism in genic regions (θ_4-fold_) was highly consistent with the pattern found in recombination rate, where significantly positive correlations were found in regions of low gene density and either no or weak negative correlation was found in regions of high gene density (Figure 5b; Table 3). After again controlling for potential confounding variables, the positive correlations remained significant in regions of low gene density among all three species (Table 3), as well as in high gene-density regions in *P. tremuloides* and *P. trichocarpa* (Table 3). We did not find any significant relationships between neutral divergence and gene density in any of the three species (Figure S14).

Compared with genic regions, correlations between intergenic diversity and gene density followed a different pattern in the three species (Figure 5c). In intergenic regions nucleotide diversity and gene density were positively correlated in regions of low gene density but negatively correlated in regions of high gene density (Figure 5c; Table 3). These correlations remained significant even after controlling for possible confounding variables (Table 3). No relationship between intergenic divergence and gene density was found in any species (Figure S14).

### Negative correlations between synonymous diversity and non-synonymous divergence at small physical scales

A negative relationship between synonymous diversity and non-synonymous divergence has been suggested to be a strong evidence of the occurrence of recurrent selective sweeps (Andolfatto 2007), and such a pattern has previously been observed in *P. tremula* using data from a small number of candidate genes (Ingvarsson 2010). Here, however, we found either no or very weak negative correlations between neutral polymorphism (θ_4-fold_) and the rate of non-synonymous substitutions (d_0-fold_) in all three species for both 100 Kbp and 1 Mbp windows, and these correlations did not change after controlling for possible confounding factors (Table 4). However, the effects of recurrent selective sweeps on synonymous nucleotide diversity are thought to be high localized within genes (Andolfatto 2007), and we therefore examined the association between θ_4-fold_ and d_0-fold_ at smaller physical scales, using data from 20,759 genes that retained more than 90% of bases after all filtering steps. In contrast to the lack of correlations observed across larger scales (100 Kbp or 1 Mbp), we found a significantly negative correlation between θ_4-fold_ and d_0-fold_ in all three species when assessed within genes (Table 4). After accounting for the possible influence of mutation rate variation among genes by normalizing θ_4-fold_ by neutral divergence rate (d_4-fold_), the negative correlations became stronger in all species (Figure S15; Table 4).

**Table 4.** Summary of the correlation coefficients (Spearman’s rank correlation coefficient) between levels of synonymous diversity (Θ_4-fold_) and non-synonymous divergence (d_0-fold_) at different physical scales in three *Populus* species.

| Dataset | Species | $d_{0\text{-fold}}$ vs. $\Theta_{4\text{-fold}}$ | |
| --- | --- | --- | --- |
|  |  | Pairwise | Partial |
| 100 Kbp<sup>a</sup> | *P. tremula* | -0.029 | -0.032 |
|  | *P. tremuloides* | -0.021 | -0.025 |
|  | *P. trichocarpa* | -0.053<sup>*</sup> | -0.051<sup>*</sup> |
| 1 Mbp<sup>a</sup> | *P. tremula* | -0.049 | 0.043 |
|  | *P. tremuloides* | -0.069 | -0.008 |
|  | *P. trichocarpa* | -0.086 | -0.006 |
| Single-genes<sup>b</sup> | *P. tremula* | -0.087<sup>***</sup> | -0.185<sup>***</sup> |
|  | *P. tremuloides* | -0.087<sup>***</sup> | -0.192<sup>***</sup> |
|  | *P. trichocarpa* | -0.148<sup>***</sup> | -0.218<sup>***</sup> |

<sup>a</sup>Partial means partial correlation controls for GC content, gene density, population recombination rate, divergence of 4-fold synonymous sites between aspen and *P. trichocarpa*, the number of 4-fold synonymous bases and 0-fold non-synonymous bases covered by sequencing data.
<sup>b</sup>Partial means correlation between $d_{0\text{-fold}}$ and $\Theta_{4\text{-fold}}/d_{4\text{-fold}}$ .
\* $P < 0.05$
\*\* $P < 0.001$
\*\*\* $P < 2.2 \times 10^{-16}$

## Discussion

### Genome-wide patterns of nucleotide polymorphism, site frequency spectrum and recombination

We have characterized and compared genome-wide patterns of nucleotide polymorphism, site frequency spectra and recombination rates in three species of *Populus*: *P. tremula*, *P. tremuloides* and *P. trichocarpa*. Although levels of nucleotide diversity varied greatly throughout the genome in all three species, we find strong genome-wide correlations of nucleotide diversity among species. It likely reflects conserved variation in mutation rates and/or shared selective constraints across the genomes in these closely related species during the time since their last common ancestor (Hudson *et al*. 1987; Charlesworth *et al*. 1993). Levels of nucleotide diversity are slightly higher in *P. tremuloides* than in *P. tremula*, and the two aspen species collectively harbor greater than two-fold levels of genome-wide diversity compared to *P. trichocarpa*. In accordance with the larger current census population size and substantially more extensive geographic range (Eckenwalder 1996), the higher genetic diversity in both aspen species most likely reflects their larger effective population sizes (*N*_e_) compared to *P. trichocarpa*. Nevertheless, interspecific variation in the mutation rate also deserves further study, particularly in light of recent results showing a feed-forward effect of genome-wide levels of heterozygosity and mutation rates (Lynch 2015; Yang *et al*. 2015). Compared to the consistent pattern of nucleotide diversity between species, the weak correlations in the allele frequency spectrum (Tajima’s D) likely reflect different demographic histories for the three species during the Quaternary ice ages (Ingvarsson 2008; Callahan *et al*. 2013; Zhou *et al*. 2014). For instance, the genome-wide excess of rare frequency alleles we observe in *P. tremuloides* is likely explained by a recent, substantial population expansion that was specific to this species.

In contrast to the mutation rate, recombination rates are only partially conserved among the three species (Figure 2c). The genome-wide average of the ratio of recombination to mutation ρ/θ_W_ or *c/μ*) was similar in *P. tremuloides* (0.39) and *P. trichocarpa* (0.38), but substantially smaller in *P. tremula* (0.22). If mutation rates are indeed unchanged between species, as suggested above, the lower estimate of *c/μ* in *P. tremula* indicates considerably lower recombination rates in *P. tremula* relative to the other two species. These discrepant results obtained from patterns of polymorphism and recombination between *P. tremula* and *P. tremuloides* likely stems from different effects of effective population size on nucleotide diversity and linkage disequilibrium (Tenesa *et al*. 2007). These effects are known to operate over different time-scales and are likely therefore differentially affected by temporal variation in the effective population size (Tenesa *et al*. 2007; Cutter *et al*. 2013). The recent population size expansion that we infer to have taken place in *P. tremuloides* can thus also explain why its recombination rate is seemingly higher than in *P. tremula*, even if they share similar levels of genome-wide polymorphism. Finally, the *c/μ* estimates we have estimated for *Populus* are in line with recent genome-wide estimates from several other plant species, such as *Medicago truncatula* (0.29) (Branca *et al*. 2011), *Mimulus guttatus* (0.8) (Hellsten *et al*. 2013) and *Eucalyptus grandis* (0.65) (Silva-Junior and Grattapaglia 2015).

### Pervasive signatures of purifying and positive selection across the Populus genome

In line with results from most other plant species (Gossmann *et al*. 2010), a majority (>50%-60%) of new amino acid altering mutations are subject to strong purifying selection (defined as *N*_e_s>10) in *Populus*. We find that the efficacy of purifying selection on weakly deleterious mutations is positively correlated with the inferred *N*_e_, with purifying selection acting more efficiently in *P. tremuloides* that has the largest *N*_e_ compared to the other two species. The same pattern is also found for rates of adaptive evolution, where estimates of the proportion of amino acid substitutions driven to fixation by positive selection are highest in *P. tremuloides* (65%), lowest in *P. trichcoarpa* (20%) and intermediate in *P. tremula* (43%). The prevalence of adaptive evolution in *Populus* contrasts markedly with the estimates in most plant species, where little evidence of widespread adaptive evolution is found (Gossmann *et al*. 2010). However, *Populus* is not unique among plants showing high rates of adaptive evolution, and similar estimates have recently been reported in both *Capsella grandiflora* (Slotte *et al*. 2010; Williamson *et al*. 2014) and a number of *Helianthus* species (Strasburg *et al*. 2011). Most earlier studies doing such estimation have been based on subsets of genes rather than genome-wide data, and more estimates from other plant species would be valuable to assess whether the high rate of adaptive evolution we find in *Populus* is widespread or exceptional.

Patterns of genomic variation contain abundant information on the relative importance of natural selection versus neutral processes in the evolutionary process (Cutter and Payseur 2013). We find that 0-fold non-synonymous sites exhibit significantly lower levels of polymorphism compared to 4-fold synonymous sites, and combined with an excess of rare variants found at 0-fold non-synonymous sites, our results suggest that the vast majority of amino acid mutations in *Populus* are under purifying selection (Larracuente *et al*. 2008). In addition, introns and 5’ UTR sites are also under some degree of selective constraint, although this constraint is much weaker than what we observe at non-synonymous sites. 3’ UTR sites seem to be either neutral or at least under comparable extents of selective constraint as 4-fold synonymous sites are (Andolfatto 2005). In contrast to genic categories, we find substantially higher levels of polymorphism in intergenic regions in all three species. Although an artifact of mapping errors due to a greater fraction of repetitive sequences in intergenic regions could not be entirely excluded, the markedly increase in diversity may also reflect either higher mutation rates or relaxed selective constraint in these regions. Future investigations are required to assess the relative contribution of these alternative factors (Kimura 1983; Begun *et al*. 2007).

Apart from strong selective constraints on protein-coding genes, multiple lines of evidence suggest that genome-wide patterns of polymorphism have been shaped by widespread natural selection in all three *Populus* species. First, we find significantly positive correlations between neutral polymorphism and population-scaled recombination rate in both genic and intergenic regions, even after controlling for confounding variables such as GC content, gene density, mutation rate and the number of covered sites by the data. While such a pattern is indicative of the action of natural selection, it could be explained by either background selection or selective sweeps. Both of these selective forces affect neutral sites through linkage, and the impact of selection on linked neutral diversity is more drastic and extensive in regions of low recombination (Begun and Aquadro 1992; McGaugh *et al*. 2012; Slotte 2014). The differences in the strength of the association between recombination and levels of neutral polymorphism likely reflect differences in the effective population size between species (Cutter and Payseur 2013; Corbett-Detig *et al*. 2015), as we observe substantially stronger signatures of linked selection in *P. tremula* and *P. tremuloides* compared to *P. trichocarpa*, matching the larger *N*_e_ inferred for these species. However, the impact of natural selection at linked sites also depends greatly on the local environment of recombination (Cutter and Payseur 2013; Slotte 2014), and in line with this we observe the strongest signatures of linked selection in *P. tremula* instead of *P. tremuloides*, consistent with the lower levels of genome-wide recombination rates we find in *P. tremula*.Different magnitudes of linked selection provide one of the major explanations for the disparate patterns of genomic variation among even closely related species (Corbett-Detig *et al*. 2015) and we find that this also holds true for the three species of *Populus* we have investigated.

Second, we find slightly negative correlations between recombination rate and the ratio of non-synonymous- to synonymous- polymorphism, but not divergence, in *P. tremula* and *P. tremuloides*, a pattern that suggests a reduced efficacy of purifying selection at eliminating weakly deleterious mutations in low recombination regions (Hill and Robertson 1966; Cutter and Choi 2010). The reduction of the efficacy of natural selection in regions of low recombination, known as Hill-Robertson interference, may help to understand patterns of partially positive correlations between gene density and recombination rate in these species (Gaut *et al*. 2007). Given the relaxed efficacy of purifying selection in regions of low recombination where weakly deleterious mutations are more likely to accumulate at a high rate, important functional elements are unlikely to cluster in these regions, as has already been shown in several other plant species (Anderson *et al*. 2006; Branca *et al*. 2011; Flowers *et al*. 2012) Consistent with this prediction (Haddrill *et al*. 2007), we find positive association between gene density and recombination rate in regions that experience low rates of recombination. In high-recombination regions where selection is more effective at eliminating slightly deleterious mutations, the association becomes much weaker in all three species. However, it remains unclear whether it is the recombination gradients that drive the functional organization of genomes in response to selection, or whether it is the gradients of functional genomic elements that in turn modify the evolution of recombination rates in *Populus*.

Third, by examining the relationship of neutral polymorphism, recombination rate and gene density, we find that levels of neutral polymorphism in genic regions are primarily driven by local rates of recombination, regardless of the density of functional genes. In contrast, we observe a more complex pattern in intergenic regions where levels of intergenic polymorphism are mainly driven by recombination rates in regions of low gene density, while in regions of high gene density levels of intergenic diversity are primarily shaped by the density of nearby genes. As we find that gene density and recombination rates co-vary in all three species, the signatures of linked selection associated with gene density could thus become obscured by rates of recombination, especially in regions of low gene density (Flowers *et al*. 2012). As shown in most plants studied so far (Nordborg *et al*. 2005; Slotte 2014), a negative relationship between gene density and levels of neutral polymorphism is more likely attributed to more intense purifying selection against deleterious mutations in regions of greater gene density, and the magnitude of such effects depends on the strength of purifying selection (Sella *et al*. 2009). In accordance with this expectation, most new mutations in genic regions are strongly deleterious and would be eliminated too quickly to remove large amounts of genetic variation at linked neutral loci. Thus even in regions of high gene density, we do not find negative correlations between gene density and genetic diversity in genic regions. However, background selection due to deleterious mutations of moderate effect in intergenic regions could account for the negative association we observe between levels of intergenic polymorphism and gene density in regions of high gene density. It is apparent that the extent to which natural selection is acting on noncoding regions of the genome in *Populus* (e.g. intergenic regions) will be an interesting avenue for future studies.

Finally, in all three *Populus* species we find significantly negative correlations between levels of synonymous polymorphism and the rate of amino acid substitution at the scale of single genes. This pattern could be driven by either recurrent selective sweeps or background selection (Charlesworth *et al*. 1993; Andolfatto 2007). However, background selection reduce local *N*_e_ due to the removal of weakly deleterious mutations, and is therefore expected to result in both reduced levels of nucleotide polymorphism and an increase of the fixation rate of slightly deleterious mutations (Charlesworth *et al*. 1993). Background selection is thus expected to affect the rates of both synonymous and non-synonymous substitutions equally, but when variation in the rates of synonymous substitution is taken into account, we find a substantially stronger (rather than weaker) negative correlation between levels of synonymous polymorphism and the rate of protein evolution. This suggest that the negative relationship we observe between non-synonymous substitution rate and levels of variation at synonymous sites is most likely driven by effects of recurrent selective sweeps in all three species (Andolfatto 2007; Sella *et al*. 2009). Furthermore, the physical scale at which these signatures of natural selection are detected carries valuable information about the strength of positive selection at the genomic level (Macpherson *et al*. 2007). Since the signatures of recurrent selective sweeps are only detectable on a genic scale, it mostly reflects relatively weak selection on the majority of adaptive amino acid substitutions and may thus explain why we do not observe the effects at either 100-Kbp or 1-Mbp scales (Macpherson *et al*. 2007; Sella *et al*. 2009).

### Conclusion and perspectives

In summary, our findings highlight multiple lines of evidence suggesting that natural selection, both due to purifying and positive selection, has shaped patterns of nucleotide polymorphism at linked neutral sites in all three *Populus* species. Compared to the predictions of the Neutral Theory which suggest that adaptations contribute negligibly to divergence between species (Kimura 1983), we find that around 20% - 65% of all amino acid substitutions are driven to fixation by adaptive evolution in *Populus*. These estimates are in accordance with the results from a number of other organisms with large effective population sizes, such as *Drosophila* (Sella *et al*. 2009), mammalian (Halligan *et al*. 2010; Carneiro *et al*. 2012) and a few plant species (Slotte *et al*. 2010; Strasburg *et al*. 2011), but substantially higher than in species with relatively small effective population sizes, such as humans and most other plant species, where little evidence of adaptive evolution has been detected (Eyre-Walker and Keightley 2009; Gossmann *et al*. 2010). Given that all three *Populus* species share similar life-cycle characteristics, such as outcrossing mating system, relatively large *N*_e_ and limited population subdivision, future studies from other long-lived forest trees are needed to investigate whether these are characteristics more generally influencing genome-wide patterns of selection in plants (Hough *et al*. 2013). Furthermore, differences in *N*_e_ and rates of recombination among the three *Populus* species are largely explaining differences in the magnitude of linked selection we observe between them.

Our analyses suggest pervasive adaptive evolution in all three species of *Populus* and although alternative hypotheses such as demographic effects could lead to spurious evidence of natural selection (Fay *et al*. 2001), the presence of linked selection could also bias inferences of demographic history (Slotte 2014). Due to the pervasive effects of linked selection we have documented in these species, our findings suggest that more attention should be paid to the process of choosing neutral sites for demographic inferences. Alternatively, new methods that allow for the joint estimation of demography and selection from genome-wide data are urgently needed.

## Acknowledgements

We are grateful to Rick Lindroth for providing access to the samples of *P. tremuloides* used in this study. We thank Carin Olofsson for extracting DNA for all samples used in this study and Robert J. Willamson for sharing a data analysis script with us. We also thank both the editor and two anonymous referees for useful comments on the manuscript. The research has been funded through grants from Vetenskapsrâdet and a Young Researcher Award from Umeâ University to PKI. JW was supported by a scholarship from the Chinese Scholarship Council.

